# Consensus molecular subtyping of colorectal cancers is influenced by goblet cell content

**DOI:** 10.1101/2020.11.17.387035

**Authors:** Samuel A. Miller, Ahmed Ghobashi, Heather M. O’Hagan

## Abstract

A critical obstacle in the field of colorectal cancer (CRC) is the establishment of precise tumor subtypes to facilitate the development of targeted therapeutic regimens. While dysregulated mucin production is a histopathological feature of multiple CRC subtypes, it is not clear how well these pathologies are associated with the proportion of goblet cells in the tumor, or whether or not this proportion is variable across all CRC. This study demonstrates that consensus molecular subtype 3 (CMS3) CRC tumors and cell lines are enriched for the expression of goblet cell marker genes. Further, the proportion of goblet cells in the tumor is associated with the probability of CMS3 subtype assignment and these CMS3 subtype tumors are mutually exclusive from mucinous adenocarcinoma pathologies. This study provides proof of principle for the use of machine learning classification systems to subtype tumors based on cellular content, and provides further context regarding the features weighing CMS3 subtype assignment.

## Introduction

Numerous histological subtypes in colorectal cancer (CRC) are defined by mucin-related phenotypes. For example, tumors with greater than 50% volume secreted mucin, are subtyped as mucinous adenocarcinoma (*1*). It is understood that these pathologies arise due to aberrant processes related to the secretion of mucin. However, it is unclear how well these pathological classifications track with goblet cell content, or in other words, the proportion of goblet cells in a tumor. As a result, it is unclear whether goblet cell content is associated with mucin-related histological subtypes, even though this may be an important consideration in explaining the heterogeneity of outcomes in patients with mucinous CRC (*2*). To date, there are no effective methods in place to rapidly query goblet cell content across many tumor samples.

The consensus molecular subtypes (CMS) of CRC were established to generate standardized tumor subtypes based on tumor gene expression profiles (*3*). The subtypes are categorized as follows: highly mutated with microsatellite instability and immune activation (CMS1), epithelial, with WNT and MYC activation (CMS2), epithelial, with metabolic dysregulation (CMS3), and mesenchymal, with stromal gene expression (CMS4). Analysis of tumor-infiltrating immune and stromal cells have identified significant enrichment of leukocytes and cancer-associated fibroblasts, explaining the immune and mesenchymal gene signatures in the CMS1 and CMS4 subtypes respectively (*4, 5*). These studies suggest that CMS subtype assignments are heavily influenced by cell populations that are present in some, but not all tumors.

The majority of CRCs are moderately differentiated (*6*) and in accordance with the TCGA COAD dataset exhibit low expression of differentiated cell type markers (*7*). A proportion of the TCGA COAD CRCs retain high expression of differentiation markers, particularly for secretory, goblet and enteroendocrine (EEC) cells, indicating differentiation is maintained in some but not all tumors (*7*). This previous work further demonstrates that the mRNA expression of strong celltype marker genes can be effectively used as a surrogate for tumor cell content. It therefore follows that if goblet cell content is highly variable in CRC, we might be able to identify a subset of CRC with high goblet cell content under one of the CMS subtype classifications. In this study we examine the expression of marker genes for goblet cells, enterocytes, intestinal stem cells, enterochromaffin cells, which are the most abundant EEC cell of the large intestine (*8*), and common EEC progenitors, which were recently shown to be enriched in some CRCs (*7*), across the different CMS subtypes. Overall, the present study establishes proof of principle for using computational methods to rapidly query goblet cell content across sequenced tumors and establishes goblet cell content as an important predictor in the assignment of the CMS3 subtype.

## Methods

### TCGA Analysis

TCGA COAD datasets were accessed from Xena Browser, and counts were normalized as previously described (*7*). To establish marker genes for each cell type, human intestine marker genes were downloaded from https://panglaodb.se (stem, goblet, enterochromaffin, and enterocyte) or compiled from previously published studies, accessed on the pubmed database (transit-amplifying, and EEC progenitor) (*9*). Cell-type marker genes were ranked by mean expression in the TCGA COAD dataset, with the top 8 genes visualized using ggplot in Figure 1B-F. All other TCGA analyses were visualized using GraphPad Prism 8 software. Clinical data including pathological subtypes of each patient were accessed through TCGAbiolinks (*10*). Unless otherwise stated, statistical significance was determined by ANOVA with correction for multiple hypothesis testing where appropriate.

**Figure 1:**
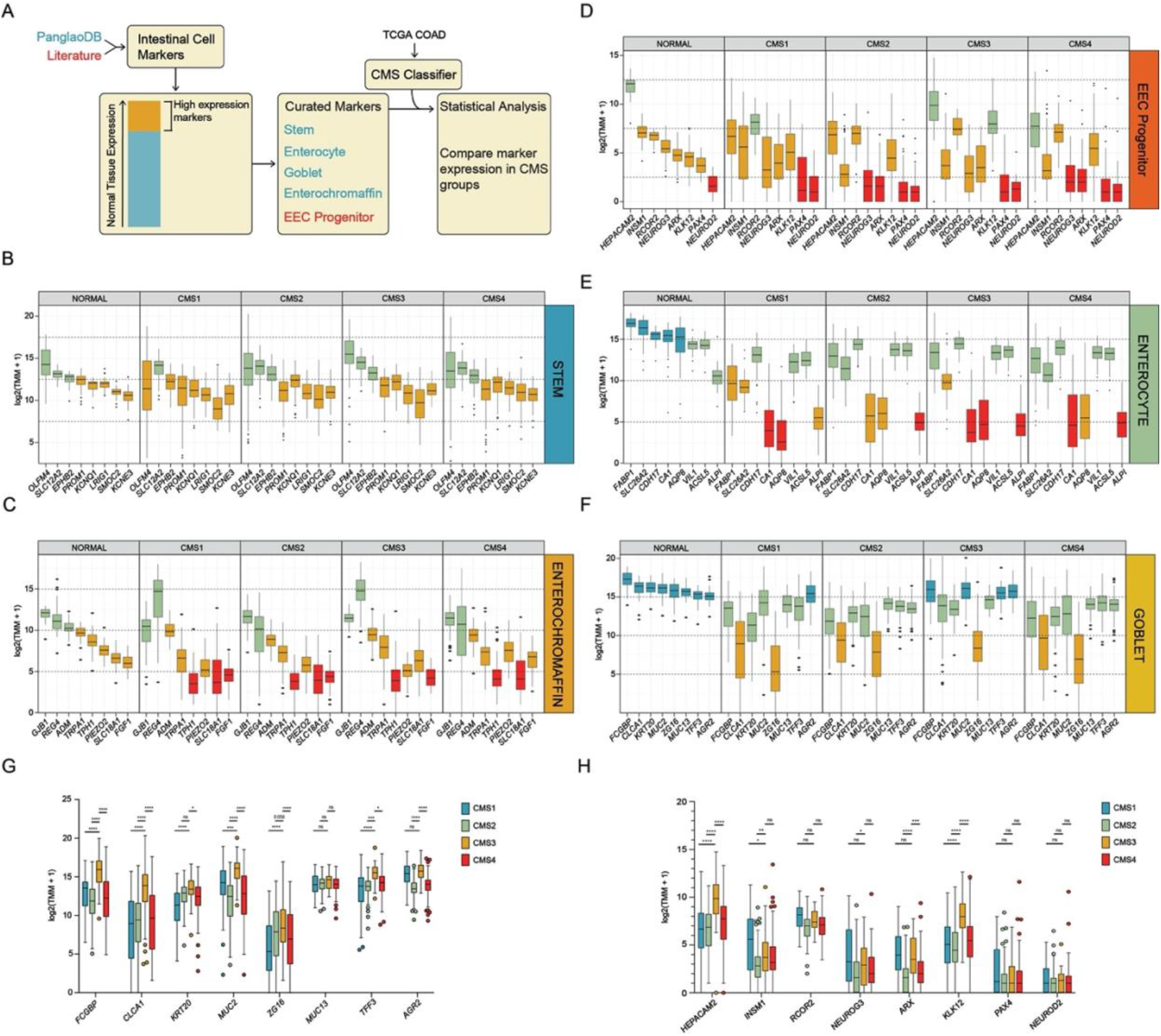
The expression levels of goblet cell markers are elevated in CMS3 subtype CRC. A, Diagram displays the order of operations in the preparation and visualization of TCGA COAD data. Blue cell type markers were acquired from PanglaoDB and red markers were curated from previously published literature accessed on PubMed. B-F, Boxplots of indicated genes expression in normal tissue and CMS subtypes: CMS1, CMS2, CMS3, and CMS4. Boxplot includes the median and box edges represent 2nd and 3rd quartiles and whiskers represent 1st and 4th quartiles, with individual dots representing outliers. Boxes are colored to clearly visualize differences in expression from high to low (blue, green, orange, red) expression respectively. Boxplots comparing (G) goblet or (H) EEC progenitor marker gene expression directly across all four CMS subtypes. Significance was determined by two-way ANOVA with pairwise Tukey’s multiple comparisons testing (*, P<0.05; **, P<0.01; ***, P<0.001; ****, P<0.0001).

### RNA extraction and qRT-PCR

The RNeasy mini kit (Qiagen, 74104) was used to extract RNA, which was then reverse transcribed into cDNA (Thermo, K1642). cDNA was amplified using gene-specific primers and FastStart Essential DNA Green Master (Roche, 06402712001). *RHOA* expression was used as a housekeeping gene for the normalization of non-housekeeping gene expression. Results are shown as mean +/- SD.

*FCGBP*, forward, AGCCACTATGAGGCGTGTTC;
*FCGBP*, reverse, CAGCCCTCATGGCATTCTGA;
*MUC2*, forward, GCTATGTCGAGGACACCCAC;
*MUC2*, reverse, AGACGACTTGGGAGGAGTTG;
*TFF3*, forward, GGAGTGCCTTGGTGTTTCAAG;
*TFF3*, reverse, AAAGCTGAGATGAACAGTGCCT;
*RHOA*, forward, CGTTAGTCCACGGTCTGGTC;
*RHOA*, reverse, ACCAGTTTCTTCCGGATGGC;

### CMS classifier

CMS classification of TCGA COAD samples was performed as previously described (*11*). Bulk RNA sequencing data of HT29 and SW480 (shEV and shLSD1) cell lines were accessed through NCBI’s Gene Expression Omnibus (GEO) series accession number GSE139927. CMS classifications of these samples were performed following log transformation using the original classifyCMS.RF function with default settings in the R package CMSclassfier (*3*).

## Results

### Goblet cell-marker gene expression is elevated in CMS3 classified colon tumors

To determine whether any particular intestinal cell types are elevated in any CMS subtype, highly expressed marker genes were plotted according to their defined CMS subtype as well as in comparison to the normal colon samples (Figure 1A). Aside from the expression of stem cell markers, all other cell type markers (enterochromaffin, EEC progenitor, enterocyte, and goblet) were generally lower in the CMS groups compared to normal colon tissue (Figure 1B-F). Additionally, differences in mean expression levels were marginal when comparing the expression of the stem, enterochromaffin, EEC progenitor, and enterocyte markers between the CMS groups. On the contrary, all goblet cell markers except for *MUC13* were generally significantly elevated in CMS3 tumors compared to other subtypes (Figure 1F, G). We have recently demonstrated that some *BRAF*-mutant CRCs are enriched for EEC progenitors and that on average, these tumors exhibit elevated expression of some goblet cell markers (*7*). Only two EEC progenitor marker genes, *HEPACAM2* and *KLK12*, were significantly elevated in CMS3 subtype tumors compared to all other CMS groups (Figure 1H). Together, these data demonstrate that CMS3 subtype CRCs exhibit higher expression of goblet cell markers. Further, CMS3 subtype CRCs do not clearly exhibit higher expression of EEC lineage progenitor or enterochromaffin markers.

### *MUC2*, but not other goblet cell markers, is elevated in mucinous CRC

The two most prevalent pathological classifications in the TCGA COAD dataset are canonical adenocarcinoma and mucinous adenocarcinoma (Figure 2A). While mucinous adenocarcinomas were uncommon in CMS2 subtype CRC (< 5%) approximately 25% of CMS1, CMS3, and CMS4 CRCs were classified as mucinous, facilitating comparison of goblet cell marker expression levels between these tumors and CMS1, CMS3, and CMS4 adenocarcinomas. *MUC2*, the major constituent of intestinal mucous (*12*), was significantly elevated in CMS1, CMS3, and CMS4 mucinous adenocarcinomas versus adenocarcinomas (Figure 2B). The only other significant comparison was the elevation of *FCGBP* expression in CMS4 mucinous adenocarcinomas versus adenocarcinomas. These data demonstrate that mucinous adenocarcinomas are not enriched in the CMS3 subtype tumors. Furthermore, mucinous adenocarcinomas are only associated with higher expression of *MUC2*, in contrast to CMS3 subtype tumors which exhibit higher expression of all goblet cell markers.

**Figure 2:**
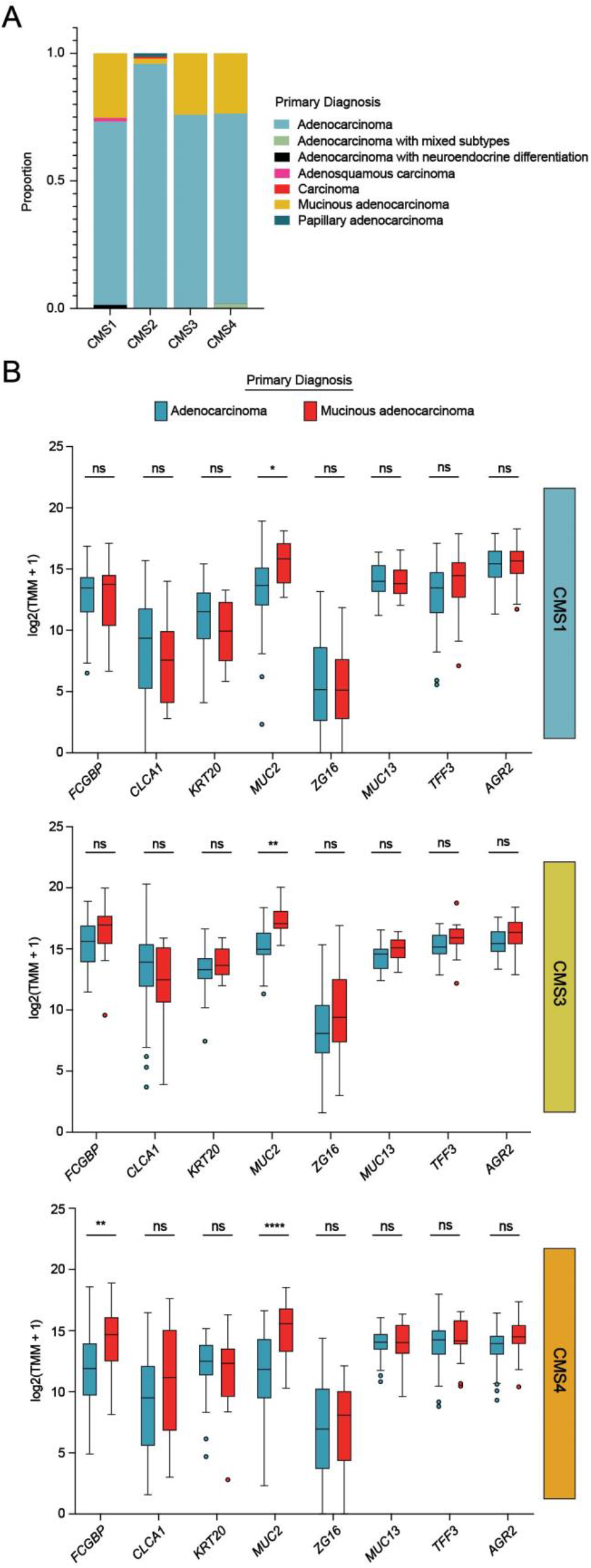
Mucinous tumors exhibit elevated *MUC2* expression but not other goblet cell markers. A, Stacked barplot depicting the proportion of each primary diagnosis in each CMS subtype from TCGA COAD. B, Boxplots of goblet cell marker gene expression subdivided by primary diagnosis (blue = Adenocarcinoma, red = mucinous Adenocarcinoma) for CMS1, CMS3, and CMS4 subtype TCGA COAD tumors. Significance was determined by two-way ANOVA with pairwise Sidak’s multiple comparisons testing (*, P<0.05; **, P<0.01; ****, P<0.0001).

### Loss of goblet cells reduces CMS3 subtype classification probability

Many CRC cell lines have previously been subtyped through the CMS classifier in an effort to define models to study each of these subtypes (*13*). The expression of goblet cell markers were analyzed in six CRC cell lines SNUC1 (CMS1), LOVO (CMS2), LS174T (CMS3), HT29 (CMS3), HCT116 (CMS4), and SW480 (CMS4) (Figure 3). In accordance with our findings in the TCGA COAD dataset, the expression of goblet cell markers *FCGBP, MUC2*, and *TFF3* were consistently expressed and elevated in CMS3 subtype cells lines, while expression was sporadic and generally orders of magnitude lower in CMS1, CMS2, or CMS4 subtype cell lines.

**Figure 3:**
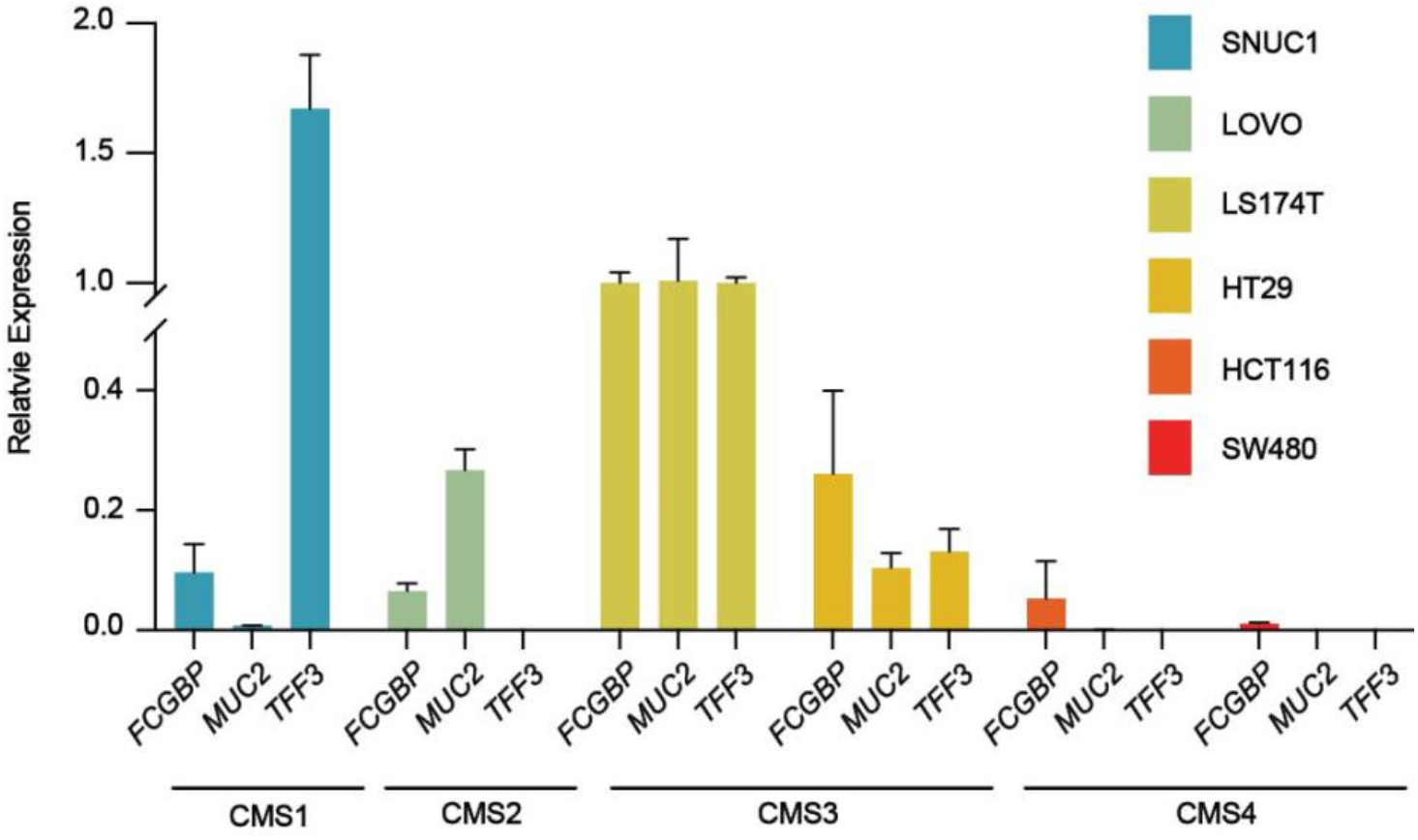
The expression levels of goblet cell markers are higher in CMS3 cell-lines compared to other subtypes. RT-qPCR of *FCGBP, MUC2, and TFF3* RNA expression level in SNUC1 (CMS1), LOVO (CMS2), LS174T (CMS3), HT29 (CMS3), HCT116 (CMS4), and SW480 (CMS4).

CMS subtyping classification relies on a series of calculations to determine the probability that any sample likely falls into any individual classification, with the assigned classification of a given tumor or cell line being the classification with the highest probability. We have demonstrated that the lysine-specific demethylase LSD1 is required for the persistence of goblet cells in the CMS3 HT29 cell line (*7*). To determine if goblet cell content is an important contributor to the probability of CMS3 subtype classification, we ran our previously published RNA-sequencing datasets comparing control empty vector (EV) versus LSD1 knockdown (KD) HT29 and SW480 cells through the CMS classifier. Consistent with previous studies, EV HT29 cells were strongly classified as CMS3 subtype in all three replicates (Figure 4A). The probability of CMS3 subtype classification was significantly reduced in LSD1 KD cells, and one replicate was now classified as having similar probability of being CMS2 or CMS3 subtype. SW480 cells are a poorly differentiated cell line that is not known to form goblet cells, consistent with low or absent expression of goblet cell markers (Figure 3). LSD1 KD did not significantly alter the probability of any CMS subtype classification in SW480 cells (Figure 4B). Together these data demonstrate that the proportion of goblet cells in a tumor is an important predictor of CMS3 subtype classification.

**Figure 4:**
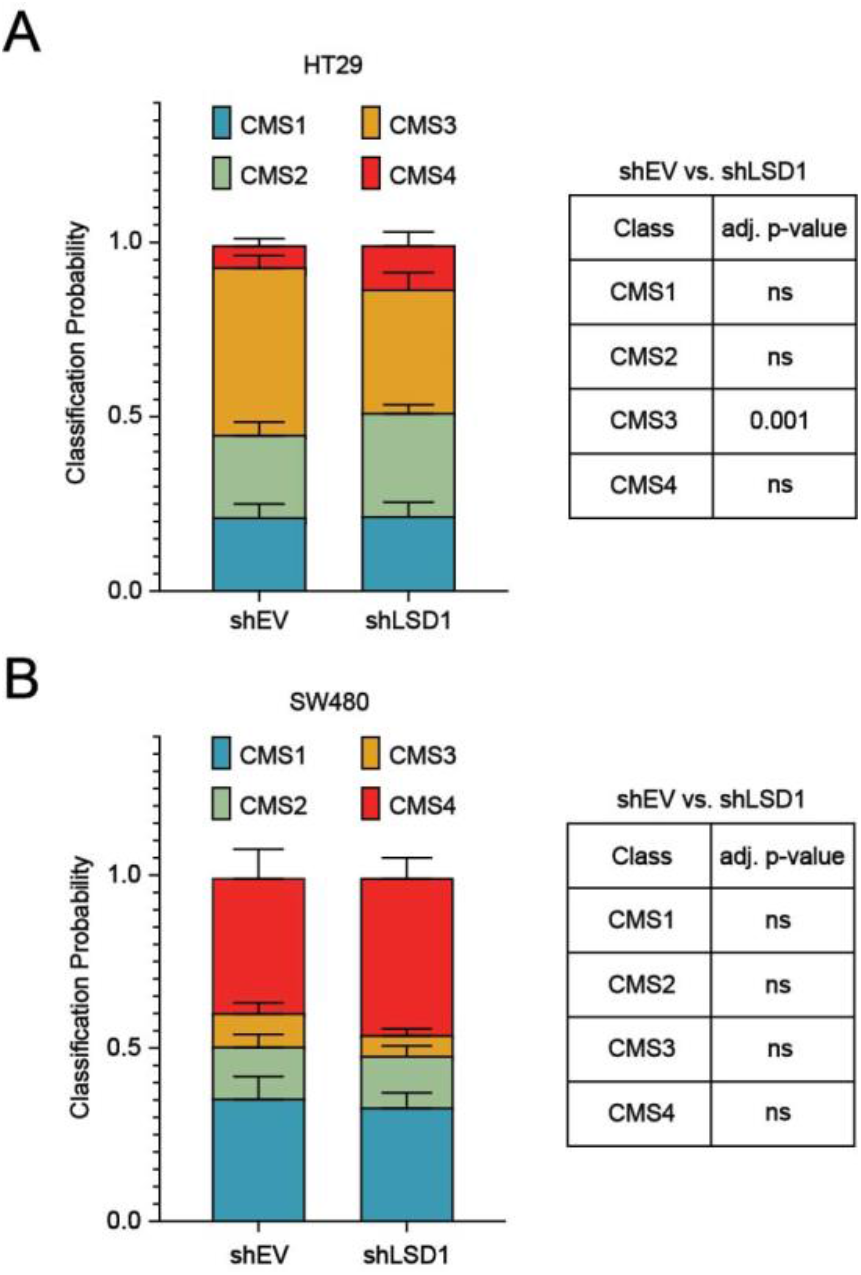
Loss of goblet cells reduces CMS3 subtype classification probability. Stacked bar plot depicting the mean probability of each CMS subtype classification +/- SD in empty vector control (shEV) or LSD1 knockdown (shLSD1) per replicate in bulk-RNA-seq experiments in (A) HT29 or (B) SW480 cells. N=3. Significance was determined by two-way ANOVA with pairwise Tukey’s multiple comparisons testing.

## Discussion

In this study, it is demonstrated that CMS3 subtype TCGA COAD exhibits elevated expression of goblet cell markers, which is an effective surrogate for goblet cell content. CMS3 subtype tumors represent a distinct subtype from COAD tumors classified as mucinous adenocarcinomas, which were only associated with high expression of *MUC2*. Analysis of CRC cell lines further confirmed that the expression of goblet cell marker genes was generally elevated in the CMS3 subtype. Lastly, analysis of RNA-seq samples from LSD1 KD, which specifically blocks the formation of goblet and early EECs, demonstrates that goblet cell content is an important factor when calculating the probability of CMS3 subtype assignment.

While markers for EEC progenitors were not generally enriched in CMS3 subtypes, *KLK12* and *HEPACAM2* were strongly elevated in the CMS3 subtype. Given the shared early lineage between EEC and goblet cells, some genes are commonly expressed in these cell types, and we have shown the EEC progenitor marker *KLK12* to be additionally expressed in goblet cells (*7*). Interestingly, mucinous adenocarcinomas showed no association with the CMS3 subtype or with the expression of goblet cell marker genes aside from *MUC2*. This study may therefore have interesting implications as it suggests that distinct mechanisms in tumors regulate the formation of goblet cells and mucin hypersecretion.

The expression of goblet cell marker genes was more consistent in the CMS3 classified compared to the non-CMS3 CRC cell lines analyzed in this study. While the expression of goblet marker genes was generally lower in non-CMS3 cell lines, SNUC1 and LOVO cells exhibited spurious high expression of one or two goblet marker genes each, which could be explained by the heterogeneous activation of transcriptional programs commonly seen in tumors (*14*). Further, although LSD1 KD significantly reduced the probability of CMS3 subtype assignment in HT29 cells, two RNA-seq replicates still exhibited slightly higher probabilities of CMS3 assignment over other subtypes. In our previous study, LSD1 KD effectively depleted goblet cells in HT29 cells, but these populations were not completely lost (*7*). The results of the CMS analysis of LSD1 KD cells suggest that residual goblet cells in this cell line may still contribute to the CMS3 subtype classification, or that in addition to goblet cell content, other factors contribute to the probability of CMS3 subtype classification.

The CMS3 subtype is substantiated by tumors with activation of various metabolic pathways (*3*). Goblet cells are highly productive, synthesizing and secreting numerous factors in addition to mucins such as trefoil factors (*15*). The goblet cell enrichment in CMS3 subtype tumors suggests that the activation of metabolic pathways in CMS3 tumors may be due to the enhanced metabolic requirements of goblet cells to produce many factors, or that the factors goblet cells secrete further activate the metabolism of non-goblet tumor cells. Most CMS subtypes are negatively correlated with one another, demonstrating clear differences in transcriptional profiles (*3*). However, this study also demonstrates that CMS2 and CMS3 exhibit almost no correlation indicating that the differences between these two epithelial subtypes may be more nuanced. We demonstrate that reducing goblet cells significantly reduces CMS3 classification in favor of CMS2 and we therefore speculate that the primary difference between these two subtypes may be the proportion of goblet cells in the tumor. Overall, this study suggests that goblet cell content may be quite variable across CRC’s. Given that the CMS classifier was able to identify tumors likely to be enriched for goblet cells, these findings support the original hypothesis that machine learning can be used as an effective means to stratify tumors based on cell content.

## References

1. F. T. Bosman, World Health Organization., International Agency for Research on Cancer., WHO classification of tumours of the digestive system. World Health Organization classification of tumours (International Agency for Research on Cancer, Lyon, ed. 4th, 2010), pp. 417 p.

2. N. I. Schneider, C. Langner, Prognostic stratification of colorectal cancer patients: current perspectives. Cancer Manag Res 6, 291–300 (2014).

3. J. Guinney et al., The consensus molecular subtypes of colorectal cancer. Nat Med 21, 1350–1356 (2015).

4. E. Becht et al., Immune and Stromal Classification of Colorectal Cancer Is Associated with Molecular Subtypes and Relevant for Precision Immunotherapy. Clin Cancer Res 22, 4057–4066 (2016).

5. P. Karpinski, J. Rossowska, M. M. Sasiadek, Immunological landscape of consensus clusters in colorectal cancer. Oncotarget 8, 105299–105311 (2017).

6. M. Fleming, S. Ravula, S. F. Tatishchev, H. L. Wang, Colorectal carcinoma: Pathologic aspects. J Gastrointest Oncol 3, 153–173 (2012).

7. S. A. Miller et al., LSD1 promotes secretory cell specification to drive BRAF mutant colorectal cancer. bioRxiv, 2020.2009.2025.313536 (2020).

8. M. L. Cristina, T. Lehy, P. Zeitoun, F. Dufougeray, Fine structural classification and comparative distribution of endocrine cells in normal human large intestine. Gastroenterology 75, 20–28 (1978).

9. O. Franzen, L. M. Gan, J. L. M. Bjorkegren, PanglaoDB: a web server for exploration of mouse and human single-cell RNA sequencing data. Database (Oxford) 2019, (2019).

10. A. Colaprico et al., TCGAbiolinks: an R/Bioconductor package for integrative analysis of TCGA data. Nucleic Acids Res 44, e71 (2016).

11. S. A. Miller et al., Lysine-Specific Demethylase 1 Mediates AKT Activity and Promotes Epithelial-to-Mesenchymal Transition in PIK3CA-Mutant Colorectal Cancer. Mol Cancer Res 18, 264–277 (2020).

12. M. E. Johansson et al., The inner of the two Muc2 mucin-dependent mucus layers in colon is devoid of bacteria. Proc Natl Acad Sci U S A 105, 15064–15069 (2008).

13. A. Sveen et al., Colorectal Cancer Consensus Molecular Subtypes Translated to Preclinical Models Uncover Potentially Targetable Cancer Cell Dependencies. Clin Cancer Res 24, 794–806 (2018).

14. L. Gonzalez-Silva, L. Quevedo, I. Varela, Tumor Functional Heterogeneity Unraveled by scRNA-seq Technologies. Trends Cancer 6, 13–19 (2020).

15. E. Aihara, K. A. Engevik, M. H. Montrose, Trefoil Factor Peptides and Gastrointestinal Function. Annu Rev Physiol 79, 357–380 (2017).

